# The stochastic logistic model with correlated carrying capacities reproduces beta-diversity metrics of microbial communities

**DOI:** 10.1101/2021.11.16.468765

**Authors:** Silvia Zaoli, Jacopo Grilli

## Abstract

The large taxonomic variability of microbial community composition is a consequence of the combination of environmental variability, mediated through ecological interactions, and stochasticity. Most of the analysis aiming to infer the biological factors determining this difference in community structure start by quantifying how much communities are similar in their composition, trough beta-diversity metrics. The central role that these metrics play in microbial ecology does not parallel with a quantitative understanding of their relationships and statistical properties. In particular, we lack a framework that reproduces the empirical statistical properties of beta-diversity metrics. Here we take a macroecological approach and introduce a model to reproduce the statistical properties of community similarity. The model is based on the statistical properties of individual communities and on a single tunable parameter, the correlation of species’ carrying capacities across communities, which sets the difference of two communities. The model reproduces quantitatively the empirical values of several commonly-used beta-diversity metrics, as well as the relationships between them. In particular, this modeling framework naturally reproduces the negative correlation between overlap and dissimilarity, which has been observed in both empirical and experimental communities and previously related to the existence of universal features of community dynamics. In this framework, such correlation naturally emerges due to the effect of random sampling.

**AUTHORS SUMMARY:** Several biological and ecological forces shape the composition of microbial communities. But also contingency and stochasticity play an important role. Comparing communities, identifying which conditions are similar in communities with similar species composition, allows to identify which forces shape their structure. Here we introduce a modeling framework which reproduces the statistical features of community similarity. We identify a single relevant parameter that captures in a single number the multidimensional nature of similarity in community composition. These results set the basis for a quantitative, and predicting, theory of the statistical properties of microbial communities.

## I. INTRODUCTION

A surprising large number of microbial species is found in a spoon of soil or a drop of water sampled at a single location and time [1]. The large values of alpha-diversity parallel with high beta-diversity: the taxonomic composition would be different if the sample were collected at a different time or in a different location [2].

A primary objective of microbial ecology is to link the observed variability of taxonomic composition with its causes. The variability of environmental conditions is one factor that determines community variability [1, 3]. Both micro-scale variability, occurring rapidly at fine spatial resolutions [4], and macro-scale variation, with spatial and temporal scale much larger than the ones of individual cells, ultimately determine which species, and in what abundance, are present at a given locations at a given time. Disturbances and environmental variability can produce an increase or decrease of beta-diversity depending on their spatial and temporal heterogeneity [5], which in turn have different effects on beta-diversity at different scales [6]. Community variability is also affected by the combination of several ecological mechanisms. For example, the immigration of a new microbial species in a community may trigger a large shift in community structure (see e.g. the case of Salmonella infection in [3], or the predictable community assembly that takes place after birth [7, 8]). Dispersal and migration are key factors in determining beta-diversity [9], as well as ecological drift and demographic stochasticity [10]. Biotic interactions and historical contingency during assembly play also an important role in shaping beta-diversity [11]. While all these factors are likely to coexist, their relative importance might vary across conditions and gradients. For instance, in plant communities, the high intra-specific aggregation in forests that produce the large values of beta-diversity are determined by different process at different latitudes [12]: environmental filtering dominates in temperate forests while dispersal limitation is more important in tropical forests.

Since the first environmental assays of the late 80s to today’s large sequencing efforts, one of the main goals of microbial communities data-analysis has been to disentangle the predictable, replicable variation of community composition — the “signal” — to contingent, non-replicable, uninformative, variability — the “noise”. Methods to identify replicable temporal or spatial patterns in the change of community composition typically rely on some measure of dissimilarity between communities — or equivalently, on a beta-diversity metric. Such a measure allows in fact to define a “distance” metric between communities, which can be ultimately used to compare and cluster communities. For example, a commonly used model-free approach is Principal Coordinate Analysis [13], which takes as input a matrix of sample-to-sample distances and identifies the coordinates which explain most of the variation between samples. This method allows to infer which variables are a more relevant source of variation and to identify clusters of similar samples. For instance, clusters identified comparing samples of microbial communities from different environments all around the world are well explained by the environment type [1]. At a smaller scale, the composition of gut microbial communities is associated with host clinical markers and lifestyle factors [14].

Despite the centrality of similarity and beta-diversity metric in microbial ecology analysis pipelines, we lack a mechanistic understanding of which aspects of community variability influence their values. Here we do not focus on disentangling the contribution of different processes in determining beta-diversity, but rather we aim at formulating a quantitative phenomenological framework able to reproduce the observed statistical properties of community similarity. The dissimilarity between two communities is in fact caused both by signal and by noise, but we miss a modeling framework that can be used to assess the effect of each. The sampling nature of the data also has a strong effect on several beta-diversity metrics [15], as it adds an additional source of noise and a bias in the observations, and should be explicitly considered.

Macroecology is a promising avenue for filling this gap. By characterising the statistical properties of community composition, macroecology provides access to quantities that are reproducible across systems. In perspective, a macroecological approach could allow to disentangle the statistical property of contingent, non-reproducible, noise from the reproducible statistical features of environmental variability.

Most of the efforts in macroecology have been focused on describing and predicting alpha-diversity. For instance, the species abundance distribution (SAD) of empirical microbiome is well characterized across ecosystems [16]. The abundance fluctuation distribution (AFD) is well described by a Gamma distribution, while the distribution of the mean abundance of species is typically well described by a Lognormal [17]. One can extend the macroecological description to dynamics, and characterise the variability of species abundance and diversity across timescales [18, 19].

One of the few examples of the study of beta-diversity under a macroecological perspective is given by the dissimilarity-overlap analysis (DOA) [20], where beta-diversity metrics have been used to infer ecological mechanisms underlying the differences in composition between samples. The DOA is based on two beta-diversity measures, dissimilarity and overlap. The dissimilarity between two communities measures the differences of the relative abundances of the species present in both samples. The overlap measures the probability that, if we pick an individual from one of the two samples, it belongs to a species that is present in both samples. These two metrics capture, in principle, two distinct aspects of community variability and should therefore vary independently. However, when they are plotted one against the other for a set of samples – in what is termed the Dissimilarity-Overlap curve (DOC)– both natural [20] and experimental [21] communities display a decreasing pattern: communities with high overlap tend to have low dissimilarity. The robustness of this DOC pattern suggests that it could be explained by a robust, general, process. The leading interpretation is that a decreasing DOC is a consequence of the universality of the dynamics [20]: different communities are subject to the same ecological dynamics, characterised by the same parameters, and differ because they occupy different dynamical attractors. Other studies have shown that other mechanism than universal dynamics might be responsible of a decreasing DOC [22]. One limit of these observations and their interpretation is that they are mostly qualitative. For instance, both the empirical DOC and models based on environmental gradients [22] produce negative DOCs. But are the empirical and modeled DOCs in quantitative agreement?

More generally, one could repeat the analysis performed by comparing overlap and dissimilarity with other beta-diversity measures. Empirical values of different beta-diversity measures are in fact correlated [23], but we lack a quantitative understanding of their relationship.

Here, we introduce a null model of community composition that is able to reproduce quantitatively the empirical values of beta-diversity and the relationship between different beta diversity metrics. The model includes two sources of variability for microbial communities. First, rapid time-variability, corresponding to the non-reproducible sources of variation. This variability was shown to be well described by the stochastic logistic model (SLM), a model that describes the temporal evolution of species abundances under a stochastic environmental noise [17, 24]. According to this model, species abundances fluctuate in time around a constant typical abundance. The second source of variability concerns how this typical abundance differs across communities [19], which represents the reproducible part of variability. The difference between these two sources, which in our model is controlled by varying a single parameter, can be of a larger or lower magnitude. Both sources of variability are modelled phenomenologically, and several mechanisms could underlie them. For instance, ecological interactions contribute to both, as they may mediate rapid abundance fluctuations and be the origin of alternative stable states in community dynamics. Environmental factors also contribute to both, either in the form of rapid environmental fluctuation or of differences in the overall environmental conditions across communities. Importantly, our model also incorporates explicitly the sampling process, allowing us to study the effect of sampling on the relationship between beta-diversity metrics and, in particular, on the DOC.

We compare the model predictions with empirical data coming from three different environments (gut, palms and oral) of different human hosts (see Methods). The model, by varying the parameter measuring the difference across-communities, jointly reproduces several beta-diversity metrics both within and across hosts. As a consequence, it also reproduces quantitatively Dissimilarity-Overlap curves, uncovering how random sampling introduces a relationship between these two in principle independent metrics.

## II. RESULTS

### A. A null model for community composition with tunable similarities

The null model for community composition that we propose is based on the statistical properties of empirical microbial communities which describe how they change across time and space. The stationary fluctuation of an OTU *i* abundance *λ*_*i*_ in time are well described by the stochastic logistic model (see [17] and Methods). Consequently, at stationarity the abundance is Gamma distributed

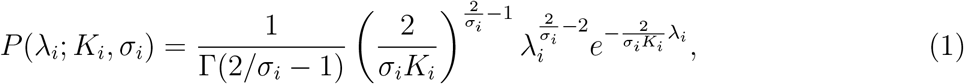

where *K*_*i*_ can be interpreted as the carrying capacity of the OTU, and *σ*_*i*_ ∈ [0, 2) is related to the coefficient of variation (see Methods). Given the compositional nature of the data, the carrying capacity of OTUs cannot be estimated from the data, as the total abundance is not known. It is however possible to estimate the values of *K*_*i*_ up to an unknown proportionality constant, which is sample-specific and common to all species in that sample [19]. For simplicity, we will still refer to *K*_*i*_ as carrying capacity in the following. With this caveat in mind, the values of *K* and *σ* for an OTU can be estimated from the time series of its abundance (see Methods). Our previous analyses [19] showed that the value of *K* for an OTU remains constant for long stretches of time. In fact, as shown in Fig. 1C, the estimations of *K* in two halves of a time series are strongly correlated. Therefore, eq. (1) together with a set of values of *K* and *σ* characterises the time-variability of composition.

**Figure 1.**
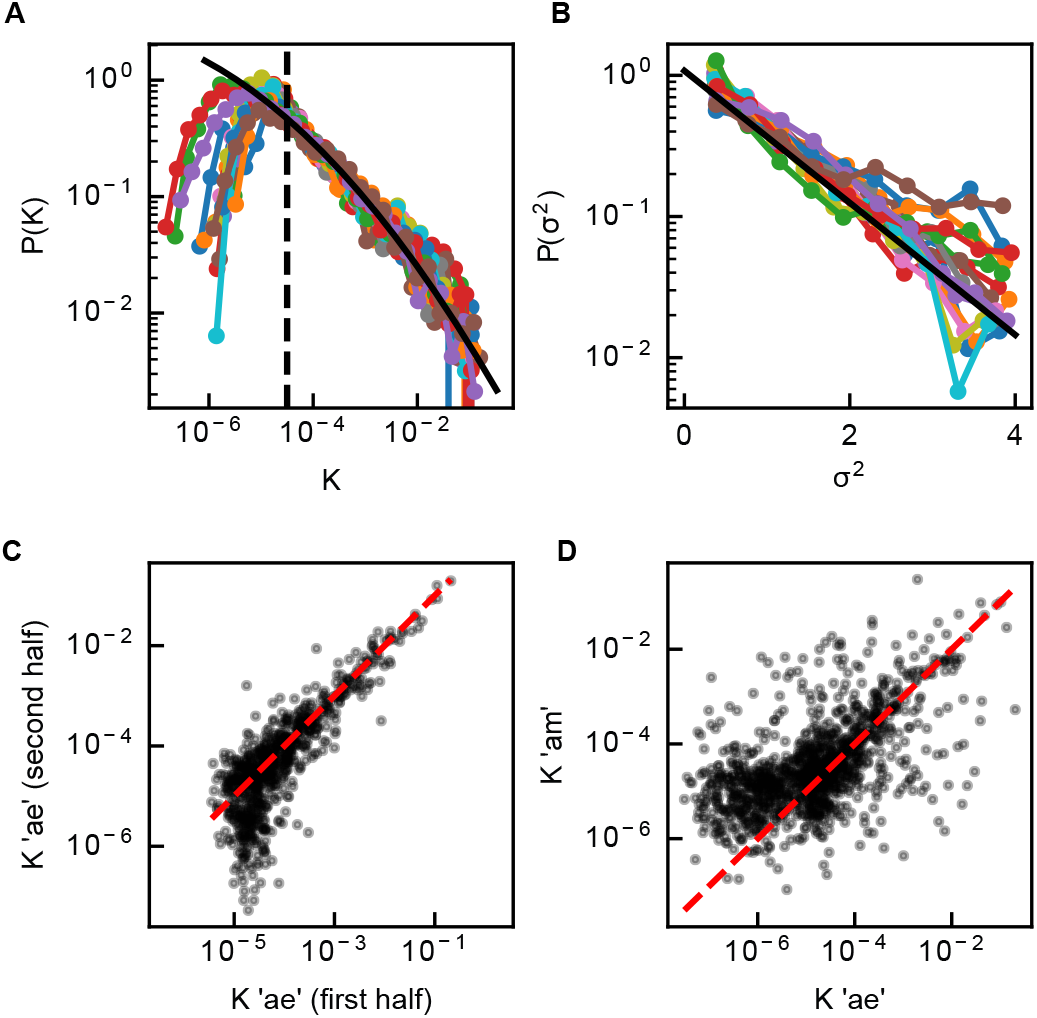
Parametrization of the null model for the gut environment. A) Distributions of *K* for each individual. The black line is a maximum likelihood fit to a truncated lognormal distribution, with truncation at 10^−4.5^, of the values from all individuals (parameters of fitted lognormal: *µ* = −19.85, *s* = 4.93); B) Distribution of *σ*^2^ for each individual. The black line is a maximum likelihood fit to an exponential distribution of the values from all individuals (mean=0.93); C) Values of *K* estimated, for each OTU of individual ‘ae’, from the first and second half of the time series. The red dotted line is the 1:1 line; D) Values of K estimated in the two individuals ‘ae’ and ‘am’, for OTUs present in both. The red dotted line is the 1:1 line.

We also compare communities inhabiting different hosts. Values of *K* estimated from the time series of two different hosts are correlated, proving that the carrying capacity is to some extent characteristic of the OTU, however the correlation is lower than the temporal one (Fig. 1D). This indicates that different communities have values of *K* that are different, although correlated. Indeed, differences in *K*, together with random fluctuations of abundance, explain almost all the dissimilarity between hosts [19]. Differences in *σ*, instead, have a much smaller role in differentiating communities.

The distribution of *K* in each of the three environments that we consider (gut, palms and oral) is lognormal above a threshold (Figs 1A and S1, dashed lines are the threshold) [17]. The existence of a threshold is due to sampling. In fact, if *N*_*s*_ is the sampling depth, OTUs with values of *K* close to or below 1*/N*_*s*_ might not be sampled. Therefore, we fit a truncated lognormal distribution (black line in Fig. 1A and in Fig. S1) to all the *K > c*, with *c* a threshold different for the different environments. For all environments, the fitted lognormal describes well the data above the threshold.

We also explored the heterogeneity of environmental variability *σ* across species. We find that the distribution of *σ*^2^ is exponential for the gut and oral environments, with similar means (respectively, 0.93 and 0.9, Figs 1B and S1A). The palm environment has, instead, a non-exponential distribution of *σ*, possibly due to larger fluctuations induced by higher rates of immigration.

To formulate our null model, we start by observing that different communities within the same environment are characterised by the same parameters of the lognormal distribution of carrying capacities and of the exponential distributions of variability *σ*^2^. See, for example, the distributions corresponding to the gut communities of different individuals in Figs 1 A and B. Therefore, we assume that each environment (for the present analysis, gut, palms or oral) has its own values of these parameters, and *K* and *σ* are identically distributed across communities within an environment. We further assume that the total number *S* of OTUs present in a community, including the ones that are present but undetected, is the same across communities within the same environment.

Then, the aim of our null model is to generate pairs of communities from the same environment with different levels of similarity. Based on the previous observations on the variability of *K* and *σ* across communities, we assume that two communities have the same values of *σ* but different values of *K*. The correlation between the logarithms of the carrying capacities, *ρ*_*K*_, tunes the level of similarity of two communities.

To generate a pair of communities from a given environment, we generate *S* pairs of values from a bivariate gaussian distribution with parameters corresponding to that environment and with correlation *ρ*_*K*_. Each pair of values corresponds to an OTU, and represents the logarithms of its carrying capacity in the two communities. For each OTU we also extract a value of *σ*^2^, common to the two communities, from an exponential distribution with the parameter corresponding to that environment. Given the set of *K* and *σ* values for a community, we extract the real abundance *λ*_*i*_ of OTU *i* from the Gamma distribution in Eq. (1) with parameters *K*_*i*_ and *σ*_*i*_. Finally, we simulate sampling. A sample of the community with number of reads *N*_*reads*_ is obtained extracting the sampled abundances from a multinomial distribution with *N*_*reads*_ trials.

This null model can be used both to generate pairs of samples from the same community at different times, setting *ρ*_*K*_ = 1, and from different communities, setting *ρ*_*K*_ *<* 1. Samples from different communities differ because of the different average abundance of OTUs *K*, the Gamma fluctuations of abundances, and random sampling. Samples of the same community at different times, instead, are obtained using the same values of *σ* and *K*, and therefore only differ due to Gamma fluctuations and random sampling.

### B. By tuning the correlation of carrying capacity the null model predicts a wide range of values of beta-diversity

We use the null model formulated in the previous section to generate in-silico communities with a range of values of community similarities, obtained by varying the correlation between carrying capacities *ρ*_*K*_. We use this ensemble of communities to study the relationships between different beta-diversity measures.

In particular, we simulate pairs of samples from different communities and from the same community at different times, and for each pair compute seven different beta-diversity measures: Pearson Correlation, Jaccard similarity, Morisita-Horn and Bray-Curtis dissimilarities, the Dissimilarity and Overlap introduced in [20] and the dissimilairty Φ introduced in [19] (see Methods). The choice of these seven different measures is aimed at covering different types of measures, accounting for presence-absence, abundance, or both, and more or less focused on common species rather than rare ones. These different beta-diversity measures have correlated values in the synthetic communities (see Fig. 2, where the measures are all plotted against the Pearson correlation). The correlation can be positive or negative because some metrics measure similarity and other dissimilarity.

**Figure 2.**
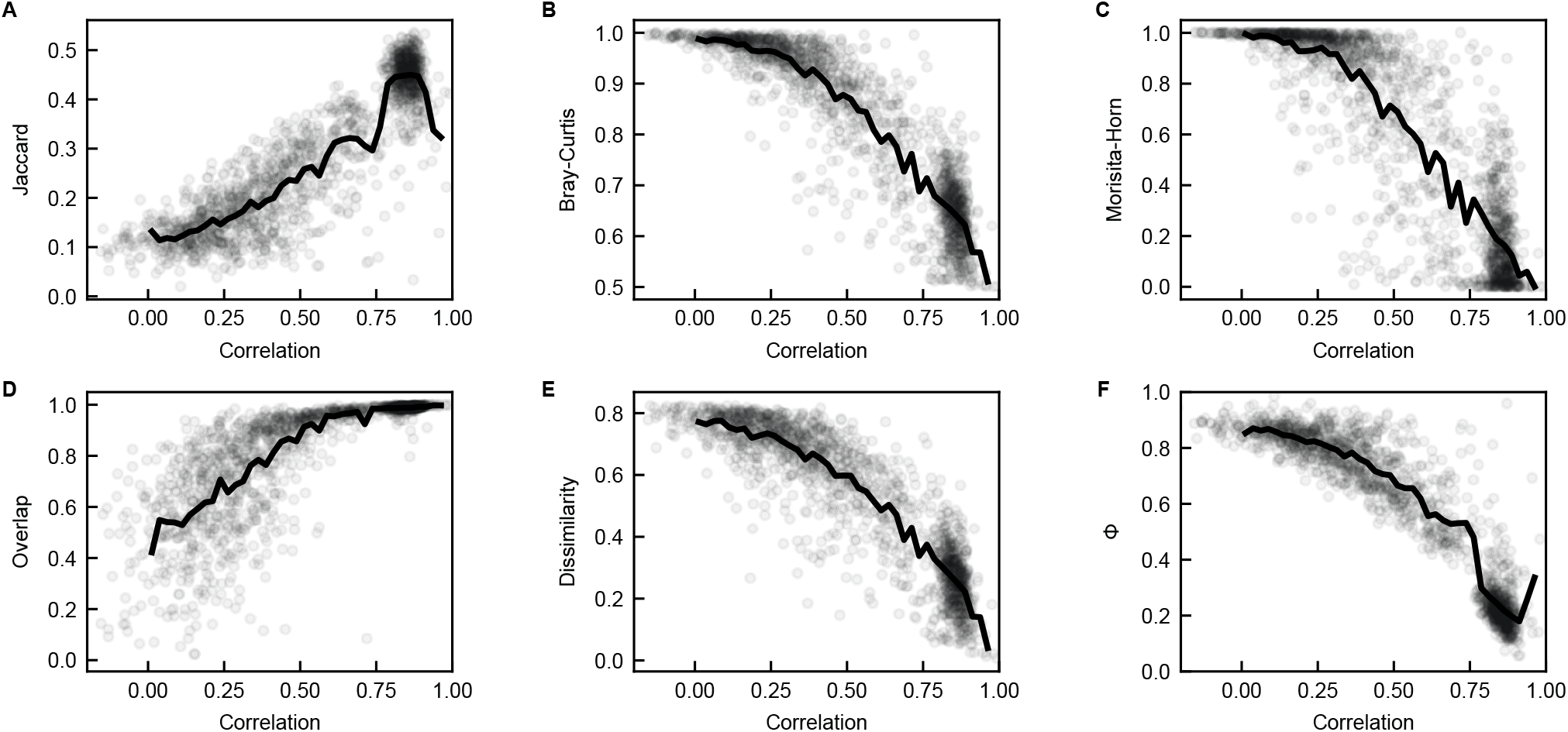
Relationships between dissimilarity measures for communities generated with the null model. The different dissimilarity measures (A: Jaccard similarity, B: Bray-Curtis dissimilarity, C: Morisita-Horn dissimilarity, D: Overlap, E: Dissimilarity, F: Φ) are plotted against Pearson Correlation. Grey circles represent the 200 pairs of communities generated with the null model. Each community has *S* = 10^4^ OTUs. Each pair of communities have the same *σ*, extracted from an exponential distribution with mean 0.9. Values of *K* are extracted from a lognormal distribution with parameters *µ* = −19 and *s* = 5. 100 pairs have the same values of *K*, to mimic samples from the same community at different times. The remaining 100 pairs have correlated values of *K*, with *ρ*_*k*_ ranging between 0.5 and 1, obtained by exponentiating values extracted from a bivariate Gaussian distribution. For each community, abundances are extracted from a Gamma distribution with parameters *K* and *σ*. For the pairs with the same values of *K*, Gamma-distributed abundances have a correlation ranging from 0 to 0.5. Reads are obtained from the real abundances by simulating multinomial sampling with number of reads 3 * 10^4^. Black lines are the binned average of the grey circles.

### C. The null model reproduces the empirical relationships between dissimilarity measures

We compare the empirical relationships between beta-diversity measures in each environment with the ones obtained from the null model with parameters fitted for that environment. Specifically, we used the fitted distributions of *K* and *σ*, a number of reads equal to the average of the reads in the empirical samples, and a number of OTUs *S* equal to the number estimated for the corresponding environment (see SI section 2). The relationships predicted by the null model follow in a remarkably precise manner the empirical patterns (Figures 3, S2 and S3). This suggests that the ingredients of the null model are sufficient to capture the statistical features of communities and of differences between communities that are relevant to determine the relationships between beta-diversity measures.

**Figure 3.**
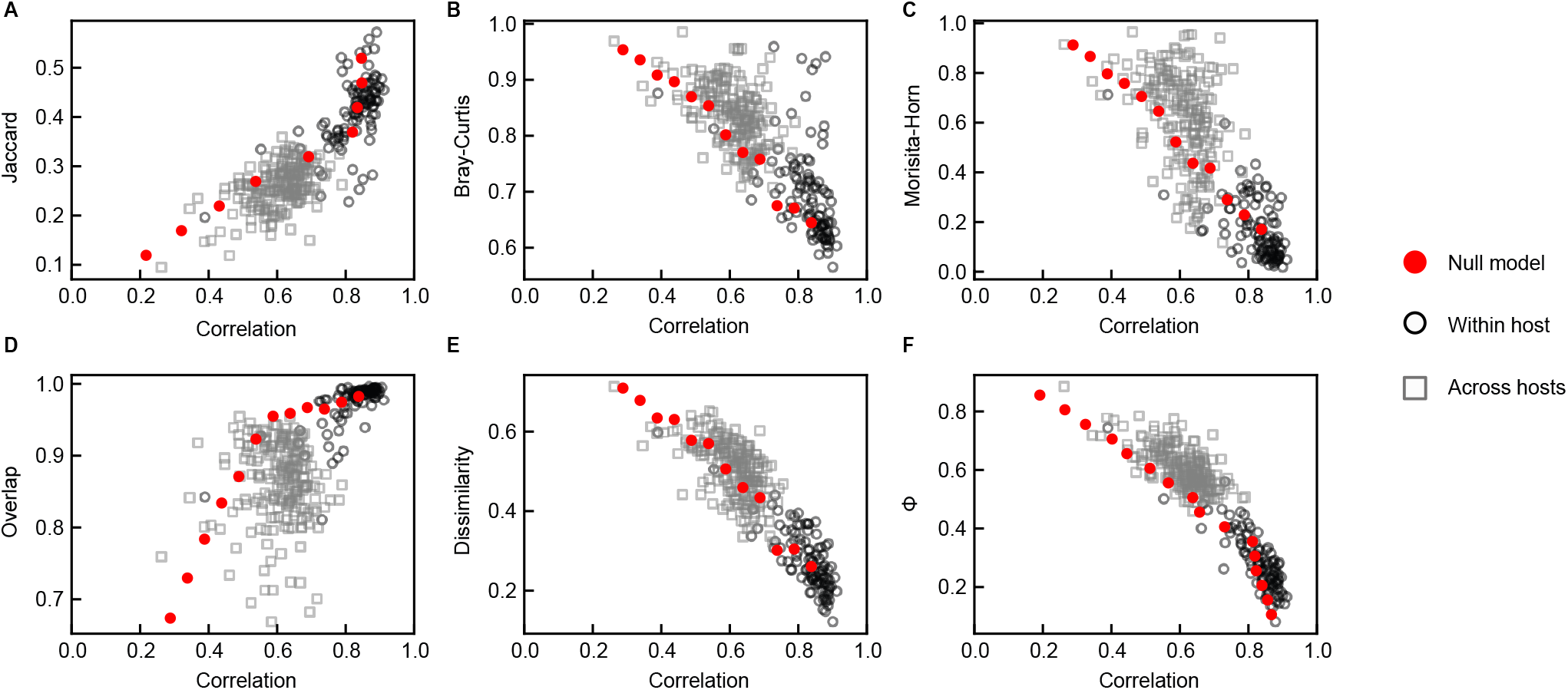
Comparison between the relationships between dissimilarity measures in empirical data for the gut environment and according to the null model. The different dissimilarity measures (A: Jaccard similarity, B: Bray-Curtis dissimilarity, C: Morisita-Horn dissimilarity, D: Overlap, E: Dissimilarity, F: Φ) are plotted against Pearson Correlation. Black circles correspond to pairs of empirical samples from the same host, while grey squares correspond to pairs of empirical samples from different hosts (but of the same dataset). Red dots are the binned average of the predictions of the null model. The null model is simulated with the distributions of *K* and *σ* fitted for the gut environment, and with a number of species equal to that estimated for the gut environment (see SI, section 2 and Table 1). The number of reads is equal to the average number of reads for the empirical samples, 3 · 10^4^.

### D. Overlap-Dissimilarity negative relationship is expected under finite sampling

Our null model shows that a decreasing Overlap-Dissimilarity curve can emerge purely because of sampling, and therefore might not reflect any ecological property of the communities.

For the communities generated with the null model, prior to sampling, overlap and dissimilarity are completely independent. In fact, all pairs of communities have overlap equal to 1, as all species are always present. However, after simulating the sampling, a non-trivial Dissimilarity-Overlap curve emerges (Fig. 4A). For pairs of communities with a medium to high correlation *ρ*_*k*_ between their (log) values of *K*, the Dissimilarity-Overlap curve has a decreasing pattern. The two insets of panel A clarify why this pattern emerges. The insets show the abundances of two pairs of communities, one with a high *ρ*_*k*_ and one with a low *ρ*_*k*_. The dissimilarity of the pair of samples is computed on the OTUs that are sampled in both, i.e. the blue points. It is clear from the insets that the abundances of the blue OTUs are more dissimilar when *ρ*_*k*_ is low. The overlap, instead, depends on the abundance of the OTUs observed in both samples (blue) with respect to that of all the observed ones (blue + orange). This quantity also has a clear pattern with *ρ*_*k*_, because the proportion of orange OTUs diminishes when *ρ*_*k*_ increases. These two effects, caused by sampling, create the the decreasing pattern in the Dissimilarity-Overlap curve.

**Figure 4.**
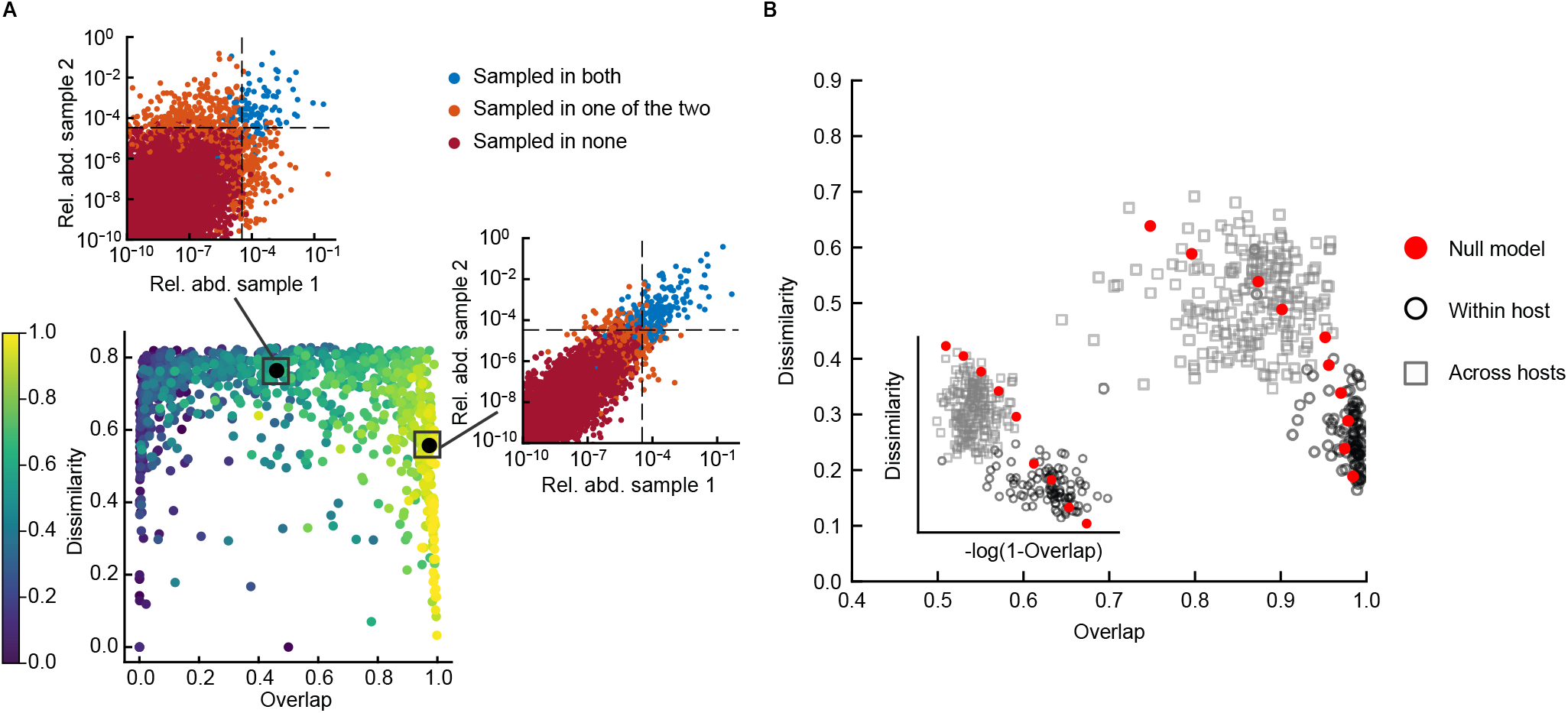
Overlap-Dissimilarity curves in the model and in empirical data. A) Relationship between Overlap and Dissimilarity for communities generated with the null model. The color of the circles corresponds to the correlation *ρ*_*k*_ of the values of *K* of the community pair, ranging from 0.6 to 1. The two insets show scatter plots of the abundances in two pairs of communities, one with a high *ρ*_*k*_ and one with a low *ρ*_*k*_. Blue circles represent OTUs sampled in both communities, orange circles OTUs sampled in only one community, and red circles OTUs sampled in neither. The dotted lines mark 1*/N*_*reads*_. B) Relationship between overlap and dissimilarity in empirical data (black circles: samples from the same hosts, grey squares: samples from different hosts) and according to the model (red circles, binned average of model prediction). The inset shows the same plot with a logarithmic scale on the x axis. For the main plot, the binned average of the model prediction is performed along the y axis, to better capture the pattern at high overlap values.

To verify if the null model can quantitatively explain the Dissimilarity-Overlap curve of empirical data, for each environment we compare the empirical pattern with that obtained from the simulation of the model with parameters measured for that environment (see explanation in the previous section). The null model is able to reproduce well the Dissimilarity-Overlap curve of empirical data (Fig. 4B for the gut environment, and Supplementary Fig. 2 for palms and oral). This result indicates that the relationship created by sampling between dissimilarity and overlap is sufficient to explain the decreasing pattern seen in empirical data.

## III. DISCUSSION

In this paper, we have introduced a modeling framework to describe the variability of community composition in space and time. The model is based on the stochastic logistic model (SLM), which describes the time variability and identifies the parameters that characterize a community, that is, the carrying capacity and the noise intensity. Our framework allows to model pairs of community with different levels of similarity by setting the correlation of their carrying capacities. The model quantitatively reproduces the values of several beta-diversity metrics, which weight in different ways the statistical properties of community composition variability. In particular the model naturally reproduces the negative relationship between overlap and dissimilarity observed in empirical [20] and experimental [21] data.

Our framework is based on several assumptions. In particular, it assumes that the SLM describes well the dynamics and properties of communities and that the difference in the abundance of species across communities can be captured by considering two SLMs with different carrying capacities. These two assumptions have been extensively studied in previous works. The SLM reproduces the dynamical properties of microbial communities as well as several macroecological patterns observed in empirical data [17, 24]. The variation of carrying capacities suffice to explain the typical difference of composition between communities [19].

A novel assumption — and result — of this work is that the differences in similarity observed across pairs of communities can be fully described and collapsed in a single parameter: the correlation between the (log) carrying capacities in the two communities, *ρ*_*k*_. The fact that, by varying this parameter, one reproduces the relationship between different beta-diversity metrics is a direct test of this assumption. In principle, other properties could matter to differentiate communities. For instance, the amplitude of abundance fluctuations *σ* could in principle differ across communities and be important to explain the observed beta-diversity. Our analysis complements the results of [19] by showing that the variability in *σ* is negligible from a macroecological perspective. Another element that could in principle determine beta-diversity is the set of species present in a community, which could differ across community. However, our model shows that differences in carrying capacity, together with finite sampling, are sufficient to explain empirical beta-diversity without the need to assume that different communities have different presence-absence patterns.

An interesting aspect of the variability *σ*, that we unveil in this paper, is that its values have a reproducible distribution across species. Interestingly, *σ*^2^ is exponentially distributed, with a similar scale parameter across individuals, in two out of three environments. We hypothesize that intense immigration could have disrupted this pattern for the ‘palms’ environment. This novel macroecological patterns adds to the list of reproducible statistical properties of microbial communities, that a comprehensive theory should be able to reproduce.

Our model does not address the biological origin of the values of the parameters and of the correlation of carrying capacities across individual. Their variation is due, potentially, to multiple biotic and abiotic factors, and to the interactions between species. Our framework clarifies that they are the effective dynamics parameters that suffice to explain the statistical properties of community composition.

Once the macroecological patterns are taken into account and the SLM is used to generate in-silico communities, one naturally reproduces the empirical values of beta-diversity metrics. This applies however only to the typical behavior: the expected value of Jaccard similarity for communities with a certain Bray-Curtis dissimilarity is very similar to the average values observed in empirical communities with the same Bray-Curtis dissimilarity. Individuals pairs of communities can deviate from this expected/average behaviour. Whether these deviations are significant, and what is their origin, is an interesting and open future direction, which, to be answered, require a null model of the kind proposed here.

Of particular relevance is the ability of our null model to reproduce the empirical Dissimilarity-Overlap curve (DOC), which raises questions on its interpretation. Bashan et al. [20] interpret the empirical Dissimilarity-Overlap curve as the consequence of the fact that species dynamics is governed by the same equations and the same parameters across communities. In this view, two communities with a similar set of species (high overlap) would have similar stable states (low dissimilarity) and vice versa, producing a DOC with negative slope. Our analysis points to an alternative origin of the empirical DOCs. For communities described by non-universal parameters (correlated but different carrying capacities), a DOC with negative slope naturally emerges due to finite sampling. Consequently, the DOC would have no implication on underlying ecological mechanisms.

Beyond the interpretation of the DOCs, our results speaks directly to the original assumptions of the dissimilatity-overlap analysis. The leading assumption behind it is that the two measures of dissimilarity and overlap are independent. This statement about independence is always defined only in the light of a null model for community composition: given a null statistical ensemble, the value of the dissimilarity is not correlated to the one of overlap. The null model considered in [20] implicitly assumes that the typical abundance of a species and its occupancy (how likely the species is to be present) are independent, which is not verified in the data. Occupancy and average abundance do in fact display a strong correlation [25], which is predicted by the SLM with finite sampling [17]. By including explicitly sampling noise, the SLM reproduces the relation between occupancy and abundance observed in the data [17] and qualifies therefore as more reasonable null model to test the independence between overlap and dissimilarity. Once the effect of sampling is included, and therefore the non-independence between abundance and occupancy is taken into account, a negative correlation between dissimilarity and overlap naturally emerges (as shown in Fig. 4).

One additional interesting prediction of our framework, is that, when samples with small enough overlap are included, the DOC curve should take the shape of an inverse U (Fig. 4). Therefore, for small enough values of the overlap, our null model predicts positive DOCs. In empirical data, the range of values of overlap is not large enough to observe this trend. In experimental data [21] it is possible to reach smaller values of overlap and an inverse-U DOC is in fact observed, as predicted by our framework.

Our results add an important step to the quantitative understanding of the structure of microbial communities. By generating more and more realistic in-silico communities, we gain an understating of what salient features of the data are the direct results of relevant biological and ecological processes, therefore allowing to disentangle general processes from contingent factors, signal from noise.

## IV. METHODS

### A. Data

We analyze human-associated microbial communities from three different environments: gut, oral and palms. For the gut environment, we consider time-series of 14 individuals coming from three different datasets: ten individuals of the BIO-ML dataset [26] (all those for which a dense long-term time-series is available), the two individuals M3 and F4 from the Moving Pictures dataset [27] and the two individuals A and B from [3]. The length of the time-series ranges from 6 months to 1.5 year, and the sampling frequency varies (daily in the most dense series). Individuals A and B from [3] both undergo a period of strong disturbance to their gut flora due, respectively, to two diarrhoea episodes during a travel abroad and a *Salmonella* infection. We exclude these periods from the analysis and consider for each individuals two separate time-series, before and after the perturbation. For the oral and palms environments, we use the time-series of the two individuals of the Moving Pictures dataset. The palms environment includes both the right and left palm. Only samples with a number of reads *N*_*reads*_ > 10^4^ are used, which completely excludes the time-series of the right palm of F4. Detail on how the raw data were analyzed can be found in the Supplementary Information of [19].

### B. Statistical properties of the fluctuations of abundances

The stationary fluctuations of OTUs abundance have been shown to follow a Gamma distribution [17]. Additionally, dynamical properties of these fluctuation suggest that abundance dynamic can be described by a Stochastic Logistic Model with environmental noise [17, 24]:

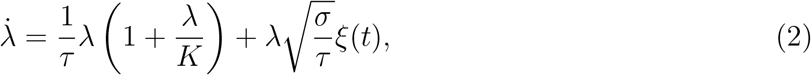

where *ξ*(*t*) is Gaussian white noise. This model has three parameters: *τ* has the dimension of a time, and determines the time-scale of relaxation to stationarity, *K* would be the carrying capacity in the absence of noise, and *σ* measures the intensity of the environmental noise. If *σ <* 2, the model has a stationary distribution which is the Gamma distribution in Eq. (1). The mean of the stationary distribution is 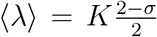 and the variance 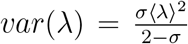. The coefficient of variation 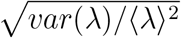 depends only on the parameter *σ*, which can thus be interpreted as the amplitude of the fluctuations. Grilli (2020) showed that Taylor’s Law applies to abundance fluctuations with exponent 2, that is, *var*(*λ*) = *C* · ⟨*λ*⟩ ^2^ with *C* a constant, which implies that *σ* is not correlated with *K*.

### C. Estimation of *K* and *σ*

The parameters *K* and *σ* can be estimated from the mean and variance of the abundance time-series, inverting the expressions for the mean and variance of the stationary distribution. To estimate the variance of abundance from the sampled abundance, we need to use an expression corrected for the sampling bias. In fact, the variance of the sampled abundance is a result of the actual variance of abundance and of the variance due to the random sampling. We use the sampling-corrected estimate as done in [17]

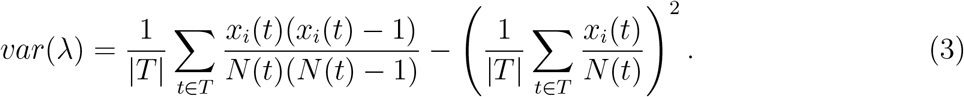

We note that the variance estimated with this formula may result negative if many counts are 0 or 1. The OTUs for which this happens are excluded from the analysis, as it is not possible to estimate their parameters.

### D. Beta-diversity measures

We consider six pairwise beta-diversity measures: Pearson correlation of log-abundances, Jaccard similarity index, the Morisita-Horn and Bray-Curtis dissimilarities, the Dissimilarity and Overlap introduced in [20] and the dissimilarity Φ introduced in [19]. The choice of these six different measures is aimed at covering different types of measures, accounting for presence-absence, abundance, or both, and more or less focused on common species rather than rare ones.

Let *n*^*A*^ and *n*^*B*^ be the OTU counts in two samples *A* and *B*, with total number of reads, respectively, *N*^*A*^ and *N*^*B*^. Let *x* = *n/N* be the sampled relative abundances. Let *S*^*A*^ and *S*^*B*^ be the sets of OTUs observed, respectively, in sample *A* and *B, S* = *S*^*A*^ ∪ *S*^*B*^ the set of all OTUs observed and *S*^*AB*^ the set of OTUs observed in both samples. Then, the beta-diversity measures are defined as follows:

#### Pearson correlation of log-abundances

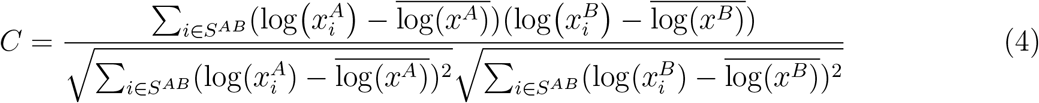

where the bar stands for an average over all *i* ∈ *S*^*AB*^. Since *x* covers several orders of magnitude, considering the logarithm allows to be sensitive to OTUs on the entire interval. Otherwise, very common OTUs would dominate the value of the correlation. However, Pearson correlation considers only OTUs observed in both samples, disregarding differences in the presence-absence pattern.

#### Jaccard similarity index

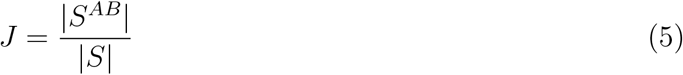

where | · | denotes the number of element of a set. The Jaccard similarity index only accounts for presence-absence, disregarding abundance. As such, it is very sensitive to differences in rare OTUs, for which a small abundance difference could cause an OTU to go undetected in a sample but not in the other.

#### Morisita-Horn dissimilarity

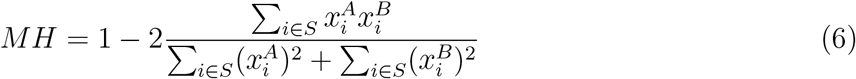

This index includes also OTUs present in only one sample (they are counted in the denominator), but is overly sensitive to common OTUs, due to the quadratic dependence on abundance.

#### Bray-Curtis dissimilarity

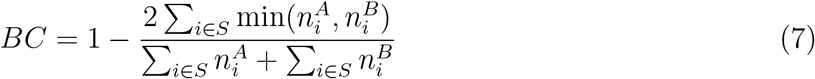

Similarly to the Morisita-Horn index, it includes also OTUs present in only one sample but is more sensitive to common ones.

#### Dissimilarity

The dissimilarity introduced in [20] is defined on the abundances 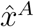 and 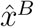, normalized on the set of OTUs common to the two samples: 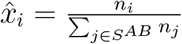. The dissimilarity is then computed as the root Jensen–Shannon divergence (rJSD) of 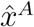 and 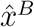:

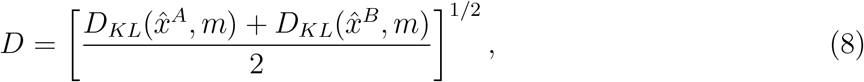

where 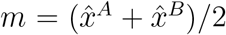 and 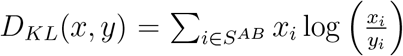 is the Kullback-Leibler divergence between *x* and *y*.

This Dissimilarity measure accounts only for the differences in abundances of OTUs common to the two samples. Additionally, the measure is dominated by common OTUs, due to the *x*_*i*_ factor in the Kullback-Leibler divergence.

#### Overlap

The overlap is the average across the two samples of the fraction of the total reads that come from OTUs observed in both samples:

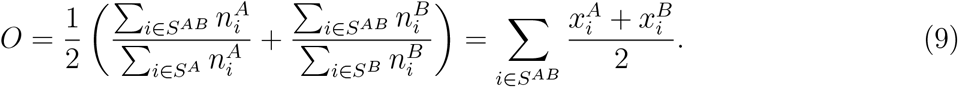

This measure accounts both for presence-absence and for abundance. In fact, two samples have a large overlap if most of the OTUs are present in both and those that are present in only one have small abundance.

#### Dissimilarity Φ

The dissimilarity measure Φ was introduced in [19] as a measure not subject to the bias due to sampling. The measure is defined on the real OTU abundances *λ* as

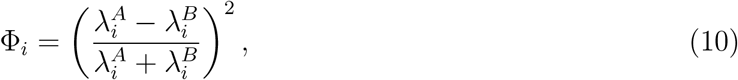

which measures the difference in abundance of OTU *i* between the two samples. Φ_*i*_ takes values in [0, 1]. It is equal to 0 when the abundances are equal and to 1 if the abundance is zero in one sample and non-zero in the other.

It is possible to show (see [19]) that its average over OTUs can be estimated from the sampled abundances. Let *ñ*^*A*^ and *ñ*^*B*^ denote the sampled abundances downsampled such that both samples have the same number of reads. Then

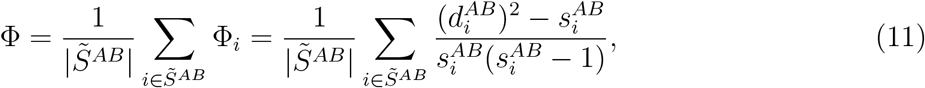

where 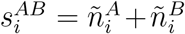 and 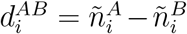, and 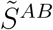 denotes the set of OTUs for which the addend is well-defined, i.e. those such that the sum of the abundances in the two samples is larger than 1.

The dissimilarity Φ gives the same importance to common and rare OTUs, as is clear from Eq. 10. Additionally, also OTUs that are only present in one sample are included in the computation (excluding those with only one count), therefore this dissimilarity measure also accounts for the presence-absence pattern.

### E. Data and code availability

This paper does not use original data. All data needed to evaluate the conclusions in the paper are available from the original references cited in the paper. The Python code used to perform the analysis is available at https://github.com/SilviaZaoli/sample_similarity.

