## Supplementary text for "The stochastic logistic model with correlated carrying capacities reproduces beta-diversity metrics of microbial communities"

#### 1 OTU selection

For the determination of the distribution of $\sigma$, we include all OTUs for which a positive $\sigma$ can be estimated from the time-series, and with an occupancy larger than 0.2. We apply this occupancy threshold to avoid an observation bias. In fact, among the very rare OTUs we only observe those with high $\sigma$ (the others are either not observed or have too low counts to estimate $\sigma$ and $K$). Therefore, including OTUs that are too rare would yield a distribution of $\sigma$ biased towards higher values. Based on previous analyses (see the Supplementary Information of [**?**]), we established the threshold at an occupancy of 0.2.

For the determination of the distribution of $K$, instead, we include all OTUs for which a positive $\sigma$ can be estimated from the time-series. In this case, in fact, the observation bias is accounted for by fitting a truncated log-normal distribution.

For all the computations of beta-diversity measures, all OTUs are included.

#### 2 Fitting a truncated log-normal distribution and estimating the total number of species

If the values of $K$ are distributed according to a probability density function $p(K)$ across OTUs, the empirical distribution of $K$ that we compute from the sampled abundances is not exactly $p(K)$. In fact, OTUs with a small $K$ might not be observed, due to the finite sampling depth. There is not a precise threshold under which OTUs are not observed. First, because the average abundance is determined not only by $K$ but also by $\sigma$, therefore two OTUs with the same $K$ could have quite different average abundances. Second, because sampling is random. However, I can establish a threshold $c$ such that OTUs with $K > c$ are almost surely observed. This threshold depends both on the sampling depth $N_{reads}$ and on the number of samples. Above this threshold, the measured $K$ will be distributed as the truncated version of $p(K)$:

$$p_{emp}(K) = \frac{\theta(K-c)p(K)}{\int dz\theta(z-c)p(z)}, \tag{1}$$

where $\theta(x-c)$ is the Heaviside function.

For our data, we observe that the measured $K$ are well-described by a log-normal distribution. Their empirical distribution above the threshold $c$ is, therefore,

$$p_{emp}(K) = \sqrt{\frac{2}{\pi s^2}}\frac{1}{K}\theta(K-c)\frac{\exp(-\frac{(\log(K)-\mu)^2}{2s^2})}{\mathrm{erfc}(\frac{log(c)-\mu}{\sqrt{2}s})}, \tag{2}$$

where $\mu$ and $s$ are the mean and variance of the normal distribution underlying the log-normal $p(K)$, that is, the mean and variance of the $\log(K)$ (of all OTUs, not just the observed ones). Instead, we call $m_1$ and $m_2$ the first two moments of the observed $\log(K)$ above the threshold:

$$m_1 = \frac{1}{S_{obs}}\sum_{i=1}^{S_{obs}}\log(K_i) \tag{3}$$

$$m_2 = \frac{1}{S_{obs}}\sum_{i=1}^{S_{obs}}\log(K_i)^2, \tag{4}$$

where $S_{obs}$ is the number of observed OTUs with $K > c$. The maximum likelihood estimate of the parameters $\mu$ and $s$ is given by the following equations:

$$m_1 = \mu + \frac{\sqrt{\frac{2}{\pi}} s \exp(-\frac{(\log(K)-\mu)^2}{2s^2})}{\text{erfc}(\frac{log(c)-\mu}{\sqrt{2}s})} \tag{5}$$

$$m_2 = s^2 + m_1\mu + \log(c)(m_1 - \mu). \tag{6}$$

After having estimated the parameters $\mu$ and $s$ for a certain environment, we can estimate the total number of OTUs $S$ for that environment starting from the number of OTUs observed above the threshold, using

$$S_{obs} = S * \int_c^\infty dK p(K) = \frac{S}{2} \text{erfc}\left(\frac{\log(c) - \mu}{\sqrt{2}s}\right), \tag{7}$$

which yields

$$S = \frac{2S_{obs}}{\text{erfc}\left(\frac{\log(c)-\mu}{\sqrt{2}s}\right)}. \tag{8}$$

For each environment, we estimate the parameters $\mu$ and $s$ by pooling together the values of $K$ above threshold from all the individuals. The value of the threshold $c$ is chosen such that none of the curves $p_{emp}(K)$ for the different individuals deviate from the fitted log-normal above $c$. Then, $S$ is estimated for each individual and averaged over individuals for the same environment, to obtain a single estimate for each environment to use in the simulations of the null model. The values of $\mu$, $s$ and $S$ for each environment are reported in Table S1.

| | Gut | Palms | Oral |
|---|---|---|---|
| $\mu$ | -19.85 | -26.73 | -16.55 |
| $s$ | 4.93 | 5.47 | 4.62 |
| $S$ | 27616 | 4253092 | 5888 |

Table S1: Estimates of the parameters of the log-normal distribution of $K$ and of the total number of OTUs for each environment.

### Supplementary figures and tables

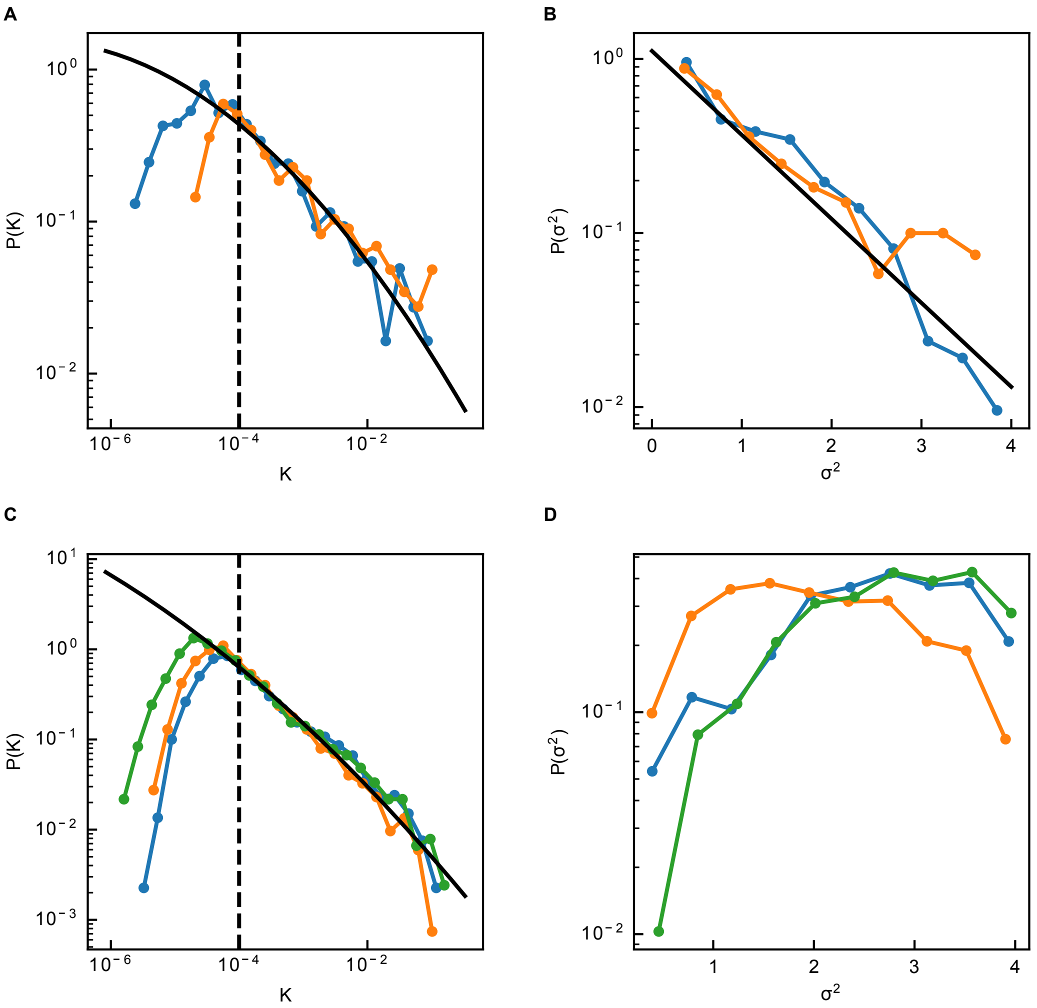


Figure S1: Parametrization of the null model for the oral (panels A and B) and palms (panels C and D) environments. A and C: Distributions of $K$ for each individual. The black line is a maximum likelihood fit to a truncated lognormal distribution, with truncation at $10^{-4}$, of the values from all individuals (parameters of fitted lognormals in Table S1); B and D) Distribution of $\sigma^2$ for each individual. In panel B, the black line is a maximum likelihood fit to an exponential distribution of the values from all individuals (mean=0.90).

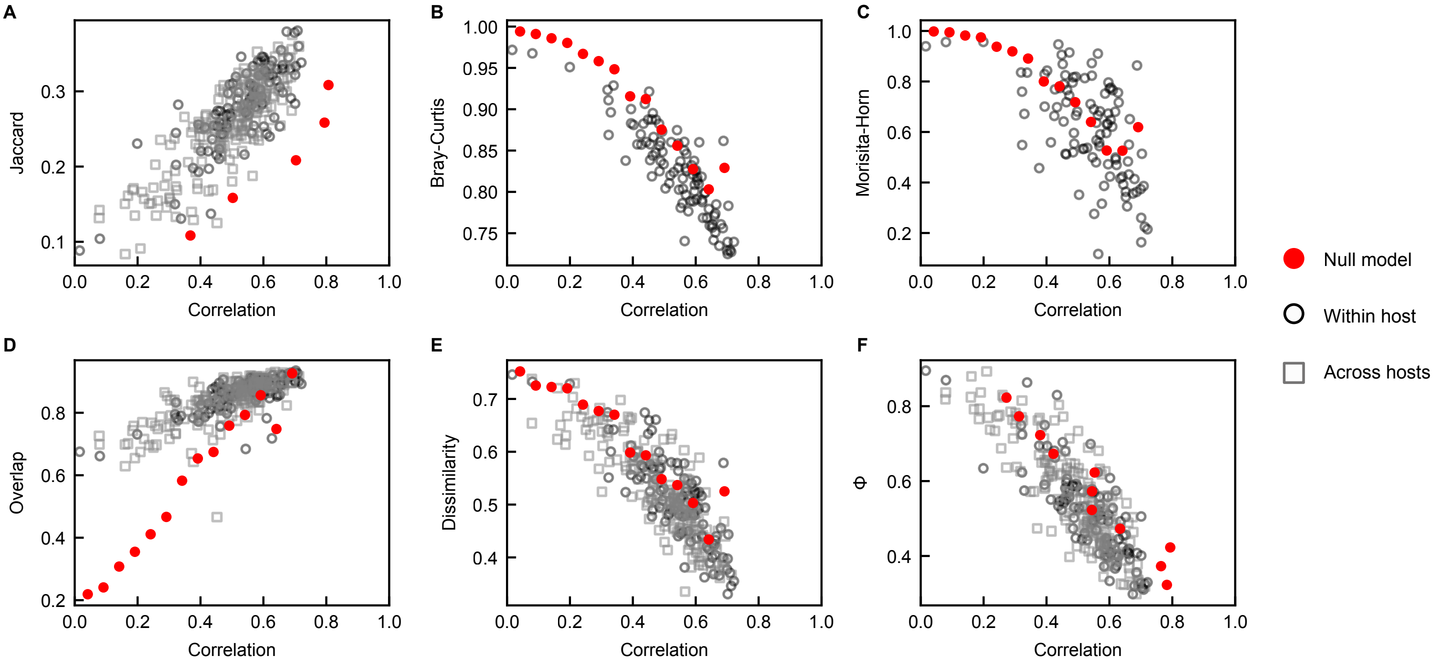


Figure S2: Comparison between the relationships between dissimilarity measures in empirical data for the palms environment and according to the null model. The different dissimilarity measures (A: Jaccard similarity, B: Bray-Curtis dissimilarity, C: Morisita-Horn dissimilarity, D: Overlap, E: Dissimilarity, F: Φ) are plotted against Pearson Correlation. Black circles correspond to pairs of empirical samples from the same host, while grey squares correspond to pairs of empirical samples from different hosts (but of the same dataset). Red dots are the binned average of the predictions of the null model. The null model is simulated with the distributions of $K$ and $\sigma$ fitted for the palms environment, and with a number of species equal to that estimated for the palms environment (see section S**??** and Table S1). The number of reads is equal to the average number of reads for the empirical samples, $2 \cdot 10^4$.

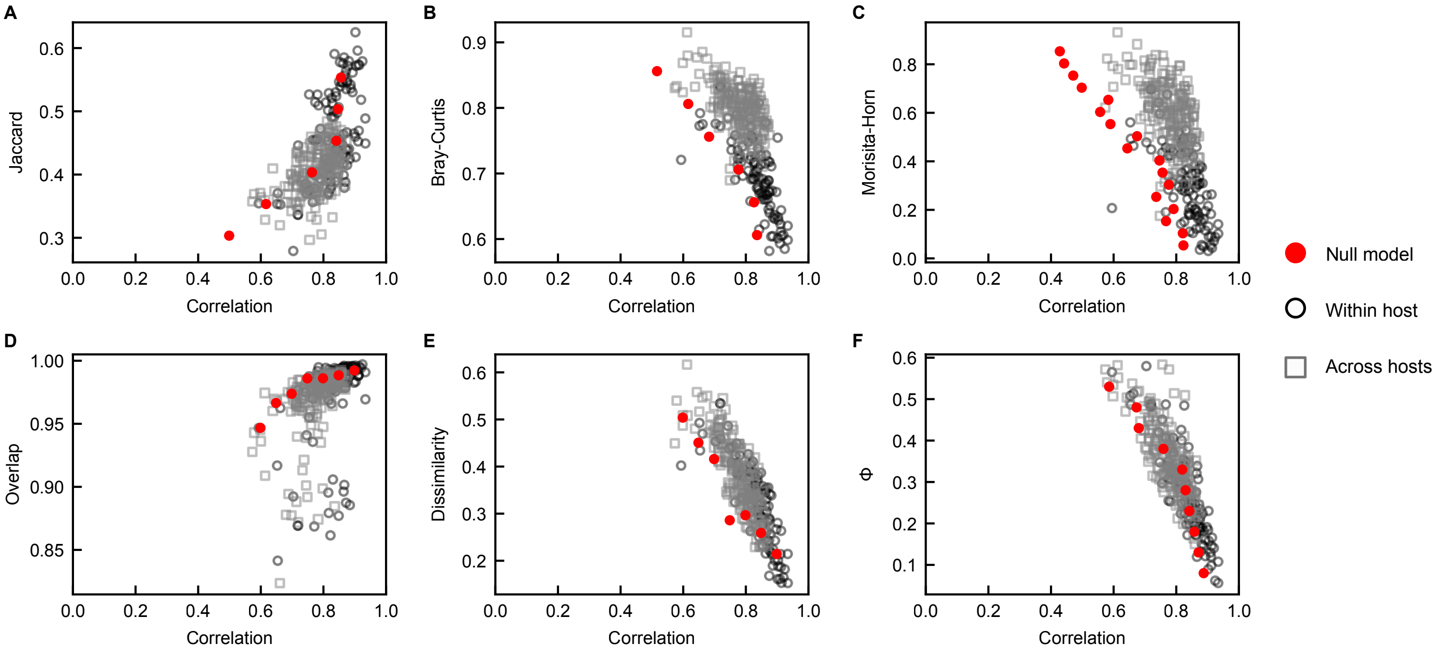


Figure S3: Comparison between the relationships between dissimilarity measures in empirical data for the oral environment and according to the null model. The different dissimilarity measures (A: Jaccard similarity, B: Bray-Curtis dissimilarity, C: Morisita-Horn dissimilarity, D: Overlap, E: Dissimilarity, F: Φ) are plotted against Pearson Correlation. Black circles correspond to pairs of empirical samples from the same host, while grey squares correspond to pairs of empirical samples from different hosts (but of the same dataset). Red dots are the binned average of the predictions of the null model. The null model is simulated with the distributions of $K$ and $\sigma$ fitted for the oral environment, and with a number of species equal to that estimated for the oral environment (see section S**??** and Table S1). The number of reads is equal to the average number of reads for the empirical samples, $3 \cdot 10^4$.

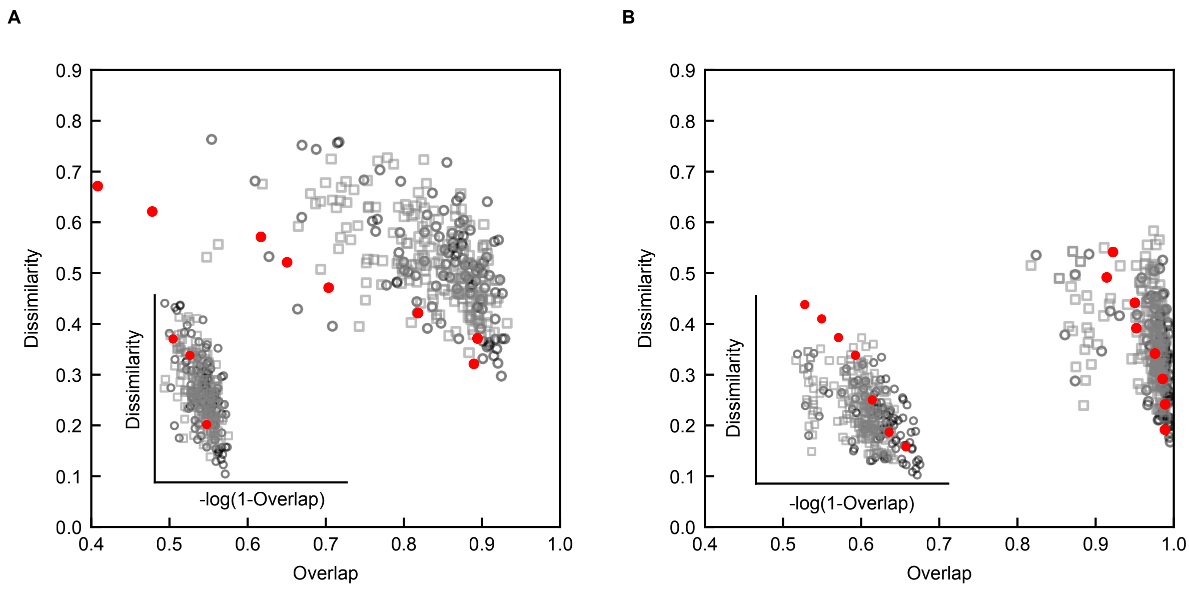


Figure S4: Comparison between Overlap-Dissimilarity curves in the model and in empirical data for the palms (A) and oral (B) environments. Black circles correspond to samples from the same host, while grey squares correspond to samples from different hosts. Red circles are binned average of model prediction. The insets shows the same plot with a logarithmic scale on the x axis. For the main plots, the binned average of the model prediction is performed along the y axis, to better capture the pattern at high overlap values.
